# Multiscale X-ray study of *Bacillus subtilis* biofilms reveals interlinked structural hierarchy and elemental heterogeneity

**DOI:** 10.1101/2021.07.27.453653

**Authors:** David N. Azulay, Oliver Spaeker, Mnar Ghrayeb, Michaela Wilsch-Bräuninger, Ernesto Scoppola, Manfred Burghammer, Ivo Zizak, Luca Bertinetti, Yael Politi, Liraz Chai

**Author notes:** To whom correspondence should be addressed: Dr. Liraz Chai, Institute of Chemistry, The Hebrew University of Jerusalem, Edmond J. Safra Campus, Jerusalem 91904, Israel. Dr. Yael Politi, B CUBE - Center for Molecular Bioengineering, Technische Universität Dresden, 01307 Dresden, Germany. **Emails:**. equal contribution (David N. Azulay, Oliver Spaeker).

## Abstract

Biofilms are multicellular microbial communities that encase themselves in an extracellular matrix (ECM) of secreted biopolymers and attach to surfaces and interfaces. Bacterial biofilms are detrimental in hospital and industrial settings, but they can be beneficial in agricultural contexts. An essential property of biofilms that grants them with increased survival relative to planktonic cells is phenotypic heterogeneity; the division of the biofilm population into functionally distinct subgroups of cells. Phenotypic heterogeneity in biofilms can be traced to the cellular level, however, the molecular structures and elemental distribution across whole biofilms as well as possible linkages between them remain unexplored. Mapping X-ray diffraction (XRD) across intact biofilms in time and space, we revealed the dominant structural features in *Bacillus subtilis* biofilms, stemming from matrix components, spores and water. By simultaneously following the X-ray fluorescence (XRF) signal of biofilms and isolated matrix components, we discovered that the ECM preferentially binds calcium ions over other metal ions, specifically, zinc, manganese and iron. These ions, remaining free to flow below macroscopic wrinkles that act as water channels, eventually accumulate and lead to sporulation. The possible link between ECM properties, regulation of metal ion distribution and sporulation across whole intact biofilms unravels the importance of molecular-level heterogeneity in shaping biofilm physiology and development.

**Significance Statement:** Biofilms are multicellular soft microbial communities that are able to colonize synthetic surfaces as well as living organisms. To survive sudden environmental changes and efficiently share their common resources, cells in a biofilm divide into subgroups with distinct functions, leading to phenotypic heterogeneity. Here, by studying intact biofilms by synchrotron X-ray diffraction and fluorescence, we revealed correlations between biofilm macroscopic architectural heterogeneity and the spatio-temporal distribution of extracellular matrix, spores, water and metal ions. Our findings demonstrate that biofilm heterogeneity is not only affected by local genetic expression and cellular differentiation, but also by passive effects resulting from the physicochemical properties of the molecules secreted by the cells, leading to differential distribution of nutrients that propagates through macroscopic length scales.

## Introduction

Biofilms are multicellular microbial communities, composed of cells that encase themselves in a secreted network of biopolymers and attach to surfaces and interfaces (1-3). Bacterial biofilms may be detrimental in hospital and industrial settings, but they are beneficial in agricultural settings. They grow on natural surfaces as diverse as rocks, plant-roots and teeth (4-6) but they also develop on synthetic surfaces such as medical devices, drinking water distribution and desalination systems (7, 8). Adopting a communal form of life, cells in biofilms organize in space and time into functionally distinct microenvironments, rendering biofilms with phenotypic heterogeneity. Phenotypic heterogeneity, the expression of specific genes in subpopulations of otherwise genetically identical cells (9-14), is related with chemical gradients arising from consumption and production processes of oxygen, nutrients, and pH changes (15), as well as local processes, such as cell death (16), presence of antibiotics (17), starvation (18), and even random activation of dedicated DNA regions (19). In recent years it has become evident that phenotypic heterogeneity serves as a survival strategy both for Gram negative and Gram positive bacteria, because it ensures the selection of at least a subpopulation in constantly changing environmental conditions (12, 19-21).

In *B. subtilis* biofilms, a model organism for biofilm formation, phenotypic heterogeneity is exhibited by the formation of subgroups of cells expressing the genes related with motility, extracellular matrix (ECM) production, sporulation (10, 22) and cannibalism (23, 24). Specifically, differential expression of ECM genes leads to regions with different mechanical properties (25-27). Importantly, as also observed in *Escherichia coli* (28), *Pseudomonas aeruginosa* (29, 30), and *Vibrio Cholerae* (31) biofilms, matrix gene expression in *B. subtilis* biofilms influences biofilm architecture, leading to local formation of wrinkles that are absent in matrix mutant biofilms (22, 25, 29, 32). In *B. subtilis* biofilms, these wrinkles act as water channels, laterally transporting water and nutrients between biofilm subgroups (33).

While phenotypic heterogeneity has been related with subgroup gene expression heterogeneity, a question that remains unanswered is whether spatio-temporal heterogeneity in biofilms is also observed at the molecular level through elemental and structural variations across the biofilms themselves. Although biochemical studies generated compositional and structural knowledge of ECM (34-40) and spore components (41, 42), these studies were mostly performed *in vitro* with isolated molecules out of the biofilm context.

To study molecular heterogeneities within intact biofilms, we used micro-focus X-ray diffraction (XRD) simultaneously with X-ray fluorescence (XRF) to scan along intact *B. subtilis* biofilm samples of different ages. XRD revealed an internal spatio-temporal division to subgroups of spores and ECM components that depended mainly on the location and hydration level of the biofilm. Simultaneous XRF measurement allowed parallel tracking of the variations in metal ion composition across biofilms. XRF measurements further exposed variable preference of biofilms, ECM components and spores to metal ions, mainly calcium, magnesium, manganese, iron and zinc. Surprisingly, the XRD signature of spores, and water as well as the XRF signals of Zn, Mn, and Fe were colocalizing at the biofilm wrinkles. These findings allowed us to portray a comprehensive multiscale molecular view of multicellularity in biofilms, demonstrating a link between biofilm morphology, molecular structure, metal ions preference and the presence of spores that mirrors the division of labor of subgroups across whole biofilms *in situ*.

## Results

### XRD of intact mature *Bacillus subtilis* biofilms reveals dominant diffraction pattern of spore- components and water molecules

Scanning XRD/XRF with a microfocused X-ray beam is a powerful tool allowing to probe the local molecular structure together with elemental mapping of heterogeneous samples with high spatial resolution. We scanned a (2.6 × 6) mm^2^ area of an intact sealed and naturally-hydrated biofilm using a micro-focused X-ray beam, as described in the schematic presented in Fig. 1. We chose a mature WT biofilm sample (10 days old) where large (mm-scale) wrinkles were preserved during sample preparation (Fig. 1). For each position along the scanned region (X-ray beam cross-section = 50×50 μm^2^, step size = 300 μm, 200 μm on X, Y axes, respectively), we simultaneously obtained a two-dimensional (2D) diffraction pattern collected on an area detector and an XRF spectrum. Simultaneous collection of XRD and XRF signals enabled correlating structural (XRD) and elemental composition (XRF) variations across a biofilm sample. The 2D XRD patterns were mostly isotropic (see for example the 2D diffraction pattern in Fig. 2A), indicating that within the biofilm plane there was no preferred alignment of structural components. We therefore azimuthally integrated the 2D patterns (in the full range 0-360°) to obtain a one-dimensional (1D) profile of the intensity (*I*) versus scattering vector (*q*). Peaks on the 1D profile correspond to reflections from ordered structures in the biofilm, with characteristic *d*-spacings (units of length) in real space inversely related to *q* (*d* = 2*π*/*q*).

**Figure 1.**
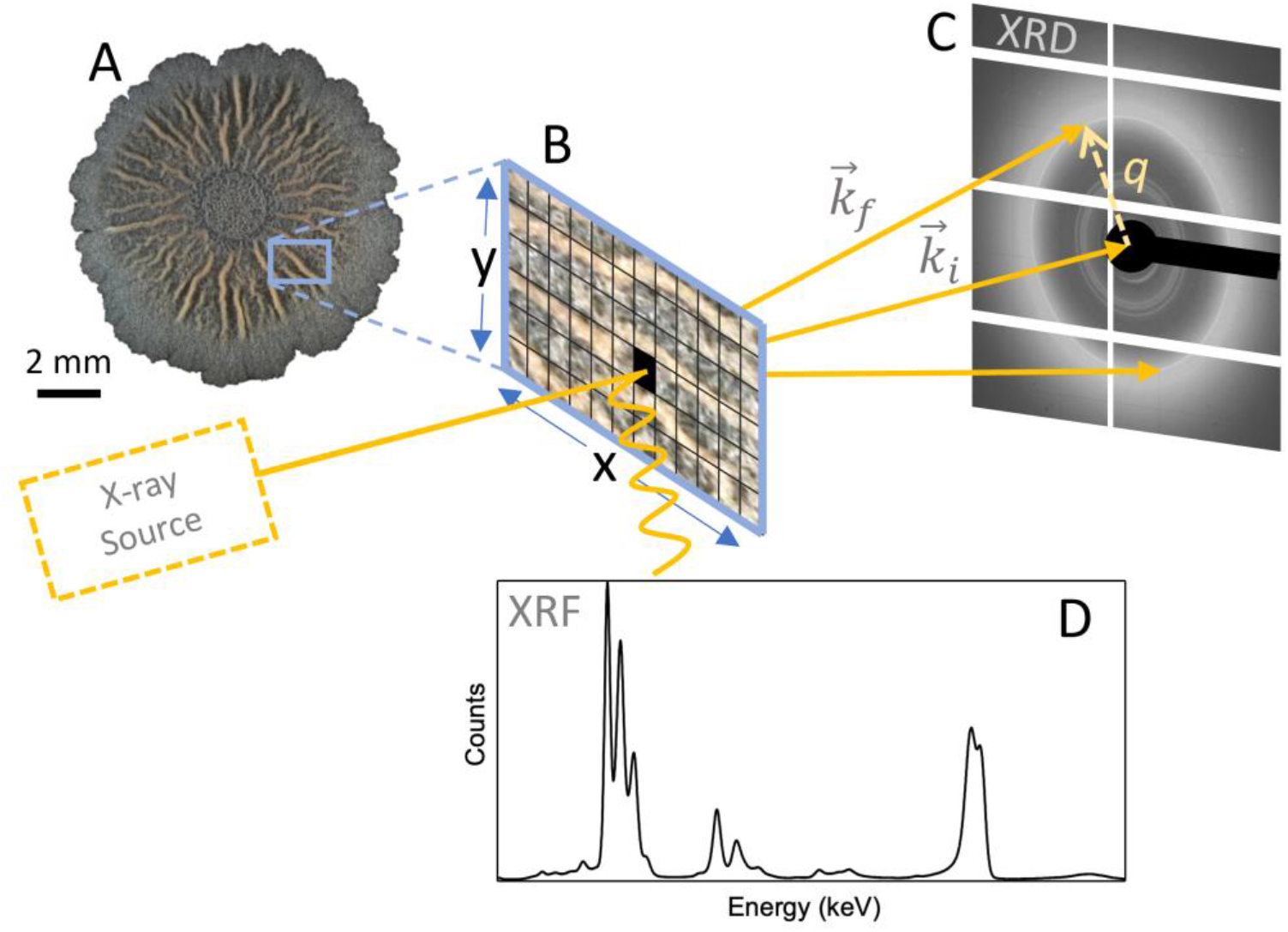
Schematic representation of the experimental set-up. A slice (B) from an intact biofilm sample (A) is placed between an X-ray source and a two-dimensional area detector (C) in transmission geometry. XRD signal is collected simultaneously with X-ray fluorescence (XRF) using a fluorescence detector (D) located almost perpendicular to the incoming beam. The X-ray beam scans the sample in pre-set step sizes along the x and y directions, defining pixels of x*y areas (see for example the black colored pixel in (B)). Scanning over large area provides structural and compositional information with spatial resolution that is determined by the size of the beam cross section and the scanning step size. 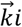 and 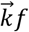 are the direct beam and scattering vectors, respectively. q is the defined as the difference between 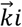 and 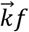. The image is not drawn to scale and the pictures used are for illustration purposes only. Random areas in the biofilm are mapped in each biofilm during an experiment.

**Fig 2.**
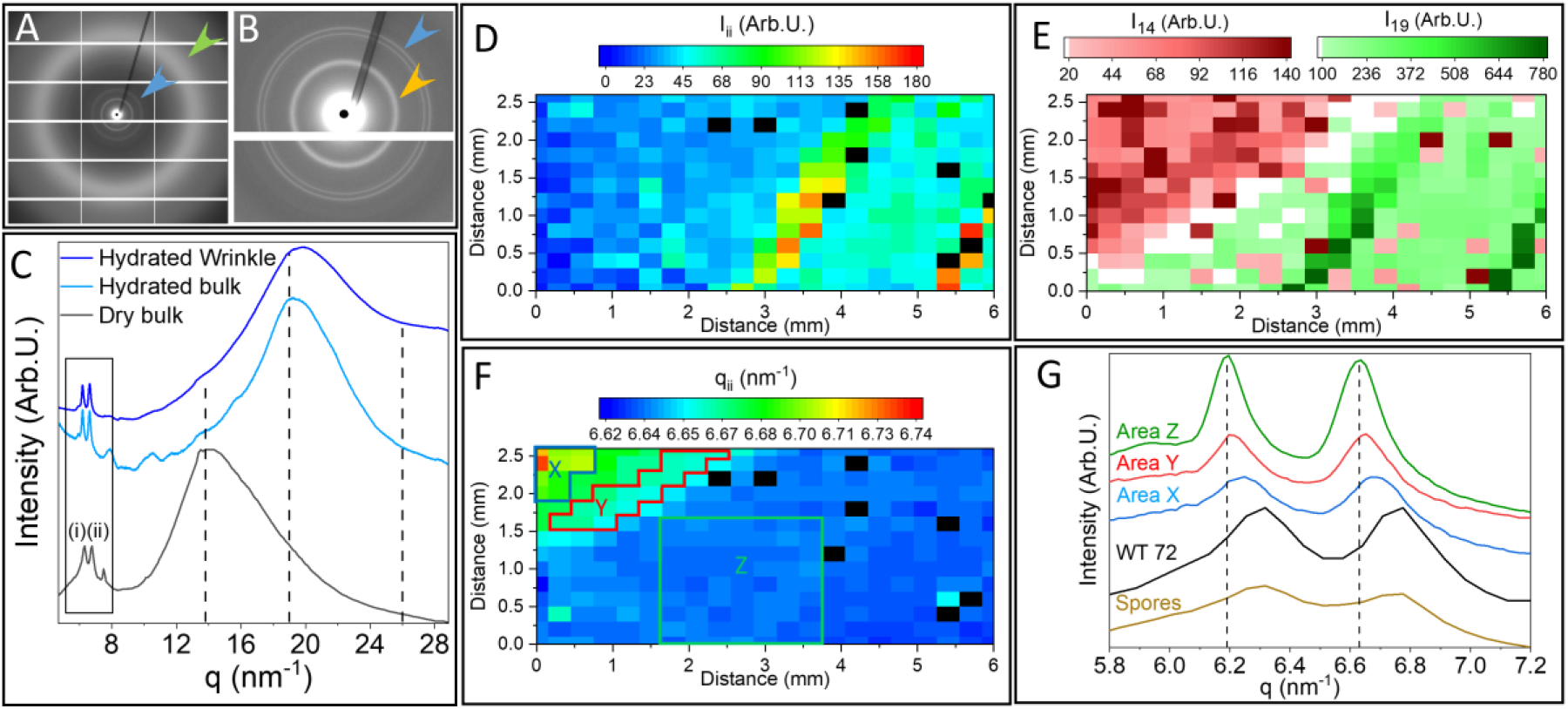
XRD of an intact hydrated and fixed-dry WT biofilm. (A) Full detector image showing a characteristic 2D pattern obtained from a natively hydrated biofilm, green arrowhead shows the water scattering signal, (B) Zoom-in into the low q range, showing the dominant doublet (marked with a blue arrowhead in A and B); Orange arrowhead marks the sample carrier (polyimide film) signal. White grid lines are gaps between detector units. (C) Representative azimuthally integrated 1D diffraction profiles of a native hydrated (dark and light blue curves taken from a wrinkle and bulk region, respectively) and fixed-dry (black curve) biofilms. The profiles show a doublet peak in the low q range (marked with rectangle and corresponding to the blue arrow in (A, B)) and broad signals in the high q range (marked with green arrowhead in (A)). Dashed lines point at peak positions q = 13.8 nm^-1^, attributed to biofilm biopolymers, and q = 19 nm^-1^, 26 nm^-1^, attributed to water scattering signal. (D) Intensity map of peak (ii) showing higher intensity along biofilm’s wrinkles. (E) Mapping the intensity of the dominant peaks in the high q range, q = 14 nm^-1^ and q = 19 nm^-1^ highlight the drying versus hydrated areas, respectively; Water scattering signal at 19 nm^-1^ is highest along the biofilm wrinkles, which serve as water channels (33). (F) Mapping of the q position of the low q peak at ∼ 6.7 nm^-1^ (peak (ii), marked in (C)), corresponding to ∼ 9.4 Å d spacing, across a WT biofilm slice of (2.6 × 6) mm^2^ area. (G) Zoom into the low q range (marked with a rectangle in (C)), showing the average profiles in different areas in the biofilm sample (X, Y, Z, in (F) corresponding to the blue, red, green curves, respectively), the average profiles in a fixed and dry biofilm (black curve) and in a spore (dry) sample (olive green curve). In fixed dry samples of 72h WT biofilms and in isolated spore samples (Fig. 2C), the d spacings were the smallest.

The diffraction patterns of the intact, hydrated, 10 days old wild-type (WT) biofilm sample showed scattering signals in both low (*q* = 5 nm^-1^ to 8 nm^-1^) and high (*q* = 10 nm^-1^ to 29 nm^-1^) *q*-ranges (Figs. 2A-C). In the low *q* range (marked with a blue arrowhead in Figs. 2A, 2B and a rectangle in Fig. 2C), WT biofilms exhibited several reflections, residing around ∼ 6 nm^-1^, corresponding to structural order with *d*-spacing of the order of 10 Å in real-space, which typically contained a sharp doublet superimposed on a broader hump (Figs. 2G, S1, Tables S1, S2). The low-q XRD pattern of dry WT biofilms largely corresponded to the XRD profile reported before for isolated spore samples (42), and confirmed by us (Figs. 2G and S1).

To evaluate the variation in specific reflections across the sample, we fitted the dominant scattering signals in the 1D XRD profiles, yielding peak position, width and amplitude in each of the measured pixels (position in the biofilm) (Fig. S1, Tables S1, S2). Mapping the spatial variation of the dominant doublet reflections in the biofilm, the doublet appeared across the whole biofilm sample, but its intensity (peak (i), Fig. S2A, and peak (ii), Fig. 2D) was highest along large wrinkles in the biofilms relative to areas away from or between the wrinkles (Fig. 2D). The relative intensities of the two peaks comprising the doublet were similar across the biofilm sample (Fig. S2C), implying that they originate from a single spore-component or from components with similar relative abundance. The doublet peak positions were rather uniform across the measured area, ranging between 6.19 to 6.24 nm^-1^ (peak (i), Fig. S2B) and 6.63-6.70 nm^-1^ (peak (ii), Fig. 2F), yet they exhibited a gradient, shifting to lower *q* values (larger d spacings) along the direction going from the top left corner of the biofilm sampling area to the bottom right (Fig. 2F).

In the high *q* range, the diffraction profiles of the hydrated sample showed typically broad peaks that were deconvoluted into three or four dominant reflections. Two reflections were fitted around 14 nm^-1^, and two reflections were centered around 19 nm^-1^ and 26 nm^-1^ (43, 44)(Fig. 2C, S4). The broad peak around 14 nm^-1^, was present in all the biofilms we tested, namely, fixed and dry (curve ‘dry bulk’ in Fig. 2C), as well as natively hydrated biofilms (‘hydrated bulk’ and ‘hydrated wrinkle’ in Fig. 2C). It includes reflections from biopolymers present in the ECM and in ordered structures inside the cells. For example, the sugar-phosphate backbone in DNA, which is present both inside and outside the cells in the biofilm (45), gives rise to a characteristic reflection at 15 nm^-1^ (46) and polysaccharides purified from biofilms showed typical scattering at *q* ∼ 13 nm^-1^ (Fig. S5).

In addition to the contribution of biopolymers to the XRD profile, native hydrated biofilms showed contributions around *q* = 19 nm^-1^ and *q* = 26 nm^-1^, corresponding to scattering from water (43, 44). In some regions, especially along wrinkles (curve ‘hydrated wrinkle’ Fig. 2C), fitting results showed that the relative amplitudes of these two peaks are comparable to those of pure liquid water, where the scattering signal originates from tetrahedral packing of water, with typical distance between oxygen atoms around 2.8 Å (44, 47) (Table S1, Fig. S4). While free water gives rise to both reflections at 19 nm^-1^ and 26 nm^-1^, bound hydration water layers in proteins lose the tetrahedral packing and as a result they give rise only to scattering signal around *q* = 19 nm^-1^ (43). Strikingly, in bulk areas away from the wrinkles (curve ‘hydrated bulk’), the free water signal at 26 nm^-1^ was diminished (Figs. 2C and S4), leading us to conclude that the contribution at q ∼ 19 nm^-1^ results from water bound to ECM and cellular biopolymers. These findings stand in agreement with a previous study, that showed that biofilm wrinkles serve as channels filled with water and essentially function to transfer nutrients across biofilms (33). In biofilms aged older than 6 days, these channels become closed tubes (33), which can explain the entrapment of free water that we observe with XRD in the 10 day old biofilm section that was removed from its original substrate.

Interestingly, the large spatial variation in the intensity of the water signal (Fig. 2E) reflects dehydration during the measurement period. Mapping the intensity of the peak contributions in the high *q* range across the sample (Fig. 2E) revealed a dehydration gradient spanning along the diagonal of the biofilm sample from the top left part of the sample, where the biopolymers reflection was dominant to the bottom right part of the sample where the water reflection at 19 nm^-1^ dominated. In the hydrated regions, the 14 nm^-1^ peak was still visible but partially obscured by the presence of the broad intense 19 nm^-1^ peak (water signal) (Fig. S4 and Table S1). The dehydration gradient provided an interesting insight into hydration/dehydration processes in biofilm spores as the drying front coincided with shifting of the spore-related doublet peak position to smaller atomic separations (larger *q* values), as shown in Figs. 2F, 2G. This strengthens the suggestion that the doublet signal is attributed to highly organized spore coat proteins (41, 42, 48, 49) and therefore sensitive to biofilm hydration. An intriguing hypothesis then arises that spore coat protein swelling may serve as humidity sensors for spore germination.

### Loss of spore-associated reflections exposes weak cross β sheet structure that is also observed in TasA fibers formed *in vitro*

The dominant contribution of the spores to the XRD signal concealed a broader hump in the low *q* region (Fig. S1). We wondered whether this hump originates from larger d spacings in ECM components, in addition to those we identified at ∼ 5 Å (corresponding to q ∼ 14 nm^-1^). Therefore, to uncover the biofilm matrix XRD signal, we performed a temporal analysis of the biofilm XRD at early time points, namely at 12 h and 24 h, where WT biofilms express matrix to a variable degree, but they do not yet produce spores (22). 1D XRD profiles of 24h old biofilms indeed showed that the spore-associated doublet, was not yet present in young biofilms (green curve, Fig. 3A, Table S1). Instead, at 24 h, broad peaks appeared at ∼ 6 nm^-1^ and ∼ 14 nm^-1^, attesting the contribution of matrix and cells to the XRD signal. At 12 h, the broad hump around 6 nm^-1^ was only occasionally present, and some regions showed only scattering signal at high q (cyan and blue curves, Fig. 3A, Table S1). Indeed, at 12 h matrix expression is initiated but the matrix is not yet uniform across the whole biofilm area (22). Probing the samples at this time point allowed us to isolate the XRD signature of cells with little contribution of the matrix. The loss of the low *q* hump in these biofilms indicated that its dependence on the presence of matrix components and that the cells themselves do not contribute to the signal at low *q*.

**Figure 3.**
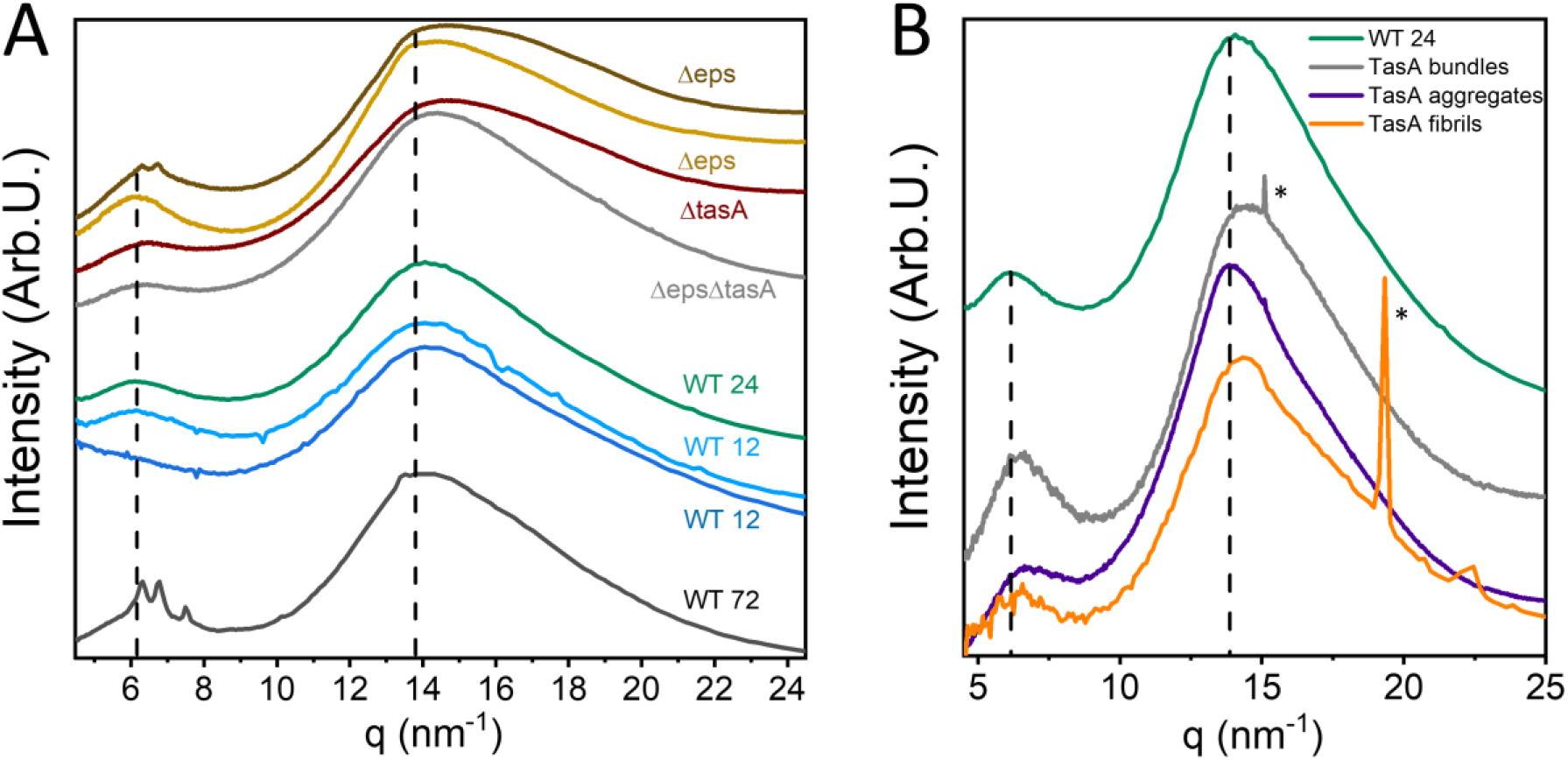
Comparison of the XRD of mature WT biofilms, young biofilms and ECM mutant biofilms. (A) XRD of fixed mature (72 h old) WT biofilms, fixed mature biofilms made by matrix mutant cells Δeps and ΔtasA, double mutant ΔepsΔtasA, as well as by fixed young biofilms (aged 12 h and 24 h), where spores have not yet developed. Dashed line marks the position of the inter-sheet and intra strand peaks in cross β sheet structures (q ∼ 6 nm-1 and 14 nm-1). (B) XRD profiles of TasA aggregates (purple curve), fibrils (orange curve), and bundles (grey curve) along with the XRD profile of 24 h old WT biofilms (green curve). The sharp peaks in the fibrils’ profile, marked with * formed at high NaCl concentration are attributed to salt crystal impurities.

We deduce that ECM contribution to the scattering signal contains a low q reflection appearing at q ∼ 6 nm^-1^ and likely an additional component at ∼ 14 nm^-1^ that reflects the atomic arrangement of the polymers. The combination of such reflections typically correspond to the canonic cross β structure attributed to a 10 Å (intrasheet) and 4.7 Å (interstrand) spacings, respectively (50).

TasA is the major matrix protein in B. subtilis biofilms and was reported to form amyloid fibers *in vitro* (34, 36, 51). We therefore also studied the XRD signal in matrix mutant biofilms, Δ*tasA*, as well as in the matrix mutant Δ*eps* that lacks the gene responsible for the production of an exopolysaccharide and the double mutant, Δ*eps*Δ*tasA* that lack both matrix components. In addition, matrix mutant biofilms were reported to negligibly express sporulation genes (22), as we also verified using Transmission electron microscopy (TEM) imaging (Fig. S6). The TEM images in Fig. S6 as well as a quantitative analysis of the cell types in different TEM images (Fig. S6D) show clearly that the abundance of sporulating cells and spores in biofilms made by the ECM mutants, Δ*eps* and Δ*tasA*, is negligible relative to their abundance in WT biofilms. The stark reduction in spore and the increased numbers of lysed cells in ECM mutant biofilms (Fig. S6), suggest increased cannibalism in matrix mutant biofilms, as previously proposed (24).

Similarly to the XRD profiles of young biofilms, the XRD profiles in *tasA* mutant biofilms (burgundy curve in Fig. 3A) as well as in the *eps* mutant biofilms (orange curve in Fig. 3A) showed loss of the spore-associated doublet. We note that some *eps* mutant biofilms also exhibited profiles where the doublet was apparent as low intensity peaks superimposed on top of a broad hump (brown curve in Fig. 3A).

XRD data of WT biofilms of different age and matrix mutant biofilms were fitted with five peak positions covering the low and high *q* ranges (see Tables S1, S2, and Fig. S1) in order to compare the dominant peak positions. Surprisingly, the cross β signature, located in the peak positions of the inter-sheet and intra strand peaks in cross β sheet structure at q ∼ 6 nm^-1^ and 14 nm^-1^ (see dashed lines in Fig. 3A), appeared in fixed mature WT biofilms and in matrix mutated biofilms (Δ*eps*, Δ*tasA* and Δ*eps*Δ*tasA*), however, the intensity of the low *q* contribution (namely around 6 nm^-1^), relative to the intensities of the high *q* contributions (Table S1), was lower in mutants that lacked TasA (burgundy curve, Fig. 3A). We conclude, therefore, that intact biofilms harbor a cross β structure that results from matrix components, and although it is not exclusive to TasA, the latter is likely the major contributor to the cross β.

Interestingly, in *eps* mutant biofilms, wrinkles are absent despite the occurrence of structure in the remaining matrix components. This observation suggests wrinkle formation depends on the co-occurrence of all the matrix components.

We were therefore intrigued to compare the structure of fibers formed *in vitro* from isolated TasA preparations with the general structural features of the ECM in intact biofilms. Recently, we have shown that TasA is polymorphic, as it forms fibers with different morphologies: aggregates in acidic solutions (termed ‘aggregates’ henceforth), fiber bundles at high protein and salt concentration (termed ‘bundles’ henceforth) and thin and long fibrils at high salt concentrations (termed ‘fibrils’ henceforth) (36, 52, 53). Fig. 3B shows the averaged 1D azimuthally integrated XRD signatures of isolated TasA that formed fibers *in vitro* under these three conditions. The XRD profiles of the three preparations of TasA fibers exhibited two broad reflections close to *q* ∼ 6 nm^-1^ and ∼ 14 nm^-1^, corresponding to the canonic cross β sheet structure. Peak-fitting shows that these peaks were slightly shifted to higher *q*, relative to those of 24 h old WT biofilms (marked by vertical dashed lines, see also Table S1), indicating a more compact packing of TasA fibers formed *in vitro* relative to biopolymers within the biofilm.

Diehl and co-workers reported XRD patterns of TasA fibers formed from recombinant protein in acidic conditions displaying narrow intra-sheet and inter-strand reflections of a cross β sheet configuration (51), which, in our case, are very broad. While the combination of these two reflections is indicative of cross β sheet features, it is unlikely that they organize into long range ordered amyloid structure in our case. This is consistent with our previous suggestions that TasA fibers form by oligomer-to-oligomer addition (53), but our data is insufficient to suggest a molecular model for the interactions between monomers and/or oligomers. We attribute the disparity in the molecular organization of TasA fibers between that reported here and that demonstrated for amyloid fibers to the fact that our fibrils are composed of full length proteins, rather than short peptides (50, 54-56) or truncated (recombinant) sequences (51). Considering that NMR studies of TasA and its crystal structure demonstrated that it contains a core of β strands arranged in a jelly roll fold, as well as alpha helices and loops (51), the cross β structure may arise from the internal core or from packing of intermolecular β strands or sheets (37). Importantly, the similarity between the XRD patterns of TasA fibers formed *in vitro* and the matrix XRD signal obtained *in situ* suggests that in intact biofilms, TasA and other ECM components form ordered fibers that are not necessarily amyloidogenic.

### Metal ions preferentially accumulate along biofilm wrinkles as well as in isolated TasA

Following the discovery that macroscopic biofilm morphology is followed by structural heterogeneity at the molecular level, we wondered whether the elemental composition of the biofilms is as heterogeneous and whether it also tracks the morphology of biofilm. Previous work of us and others have shown that metal ions accumulate in biofilms (57-59), however their spatial distribution across intact biofilms has not been addressed.

XRF signal results from the emission of X-ray photons following electron relaxation after its excitation by the X-ray beam and it is indicative of the elemental composition of a sample. The XRF signal from biofilms was recorded concurrently with XRD to probe the distribution of accumulated metal ions across biofilms (Fig. 4). The fluorescence photon counts were then converted to relative atomic density taking into account the energy dependent absorption cross-section for each element (60-63). While Ca was the most abundant metal ion in all samples, and was rather homogenously distributed across the biofilms, we found an interesting spatial distribution for Zn, Fe and Mn with increased signals along the biofilm wrinkles. A similar trend was observed for Mg (Fig. S2D), however the signal to noise in this measurement was relatively low due to the low fluorescence emission energy of Mg leading to significant absorption in air. For comparison, the distributions of Ca and K (Fig. 4, Fig. S2E, respectively) did not correlate with the wrinkle morphology.

**Figure 4.**
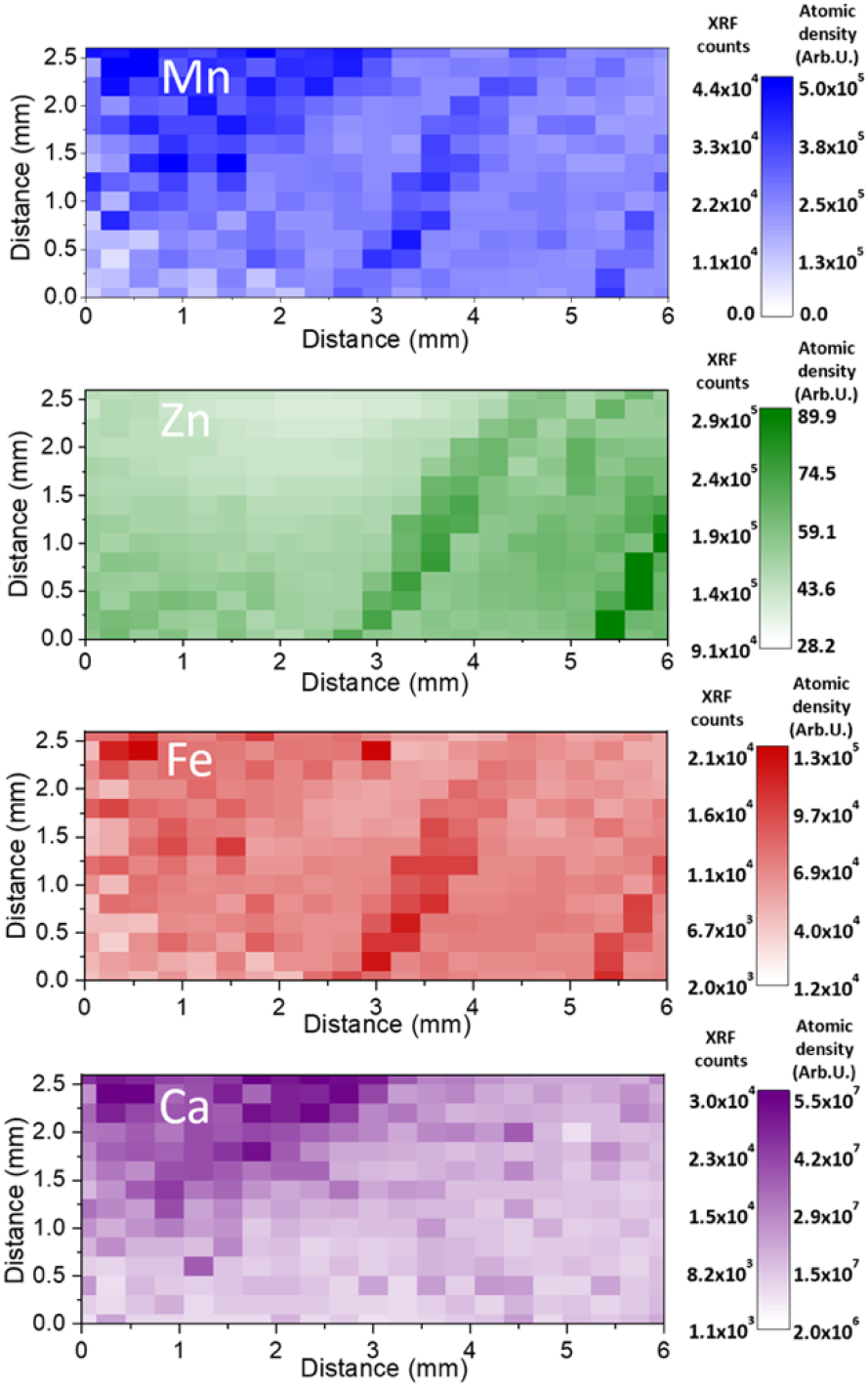
Atomic density measured by XRF of intact hydrated WT biofilm slice. XRF/Elemental distribution maps are plotted for each metal ion. The LUT describes the XRF counts (values to the left of the scale) and the corrected fluorescence signals (values to the right of the scale) of Mn (blue), Zn (green), Fe (red), Ca (magenta) of WT biofilms. The signal of Zn, Mn, and Fe, but not Ca, is stronger in the biofilm wrinkles, coinciding with water and spore accumulation along biofilm wrinkles.

The elevated signal of Zn, Mn, and Fe coincided with increased intensity of the spore-related doublet XRD signature and water scattering acquired simultaneously. A possible explanation to the increased XRD signal along the wrinkles could be that the biofilm is thicker along wrinkles, however, this effect alone would lead to an increase of the XRF signal of all the metal ions. The uniform Ca and K signals across the biofilms, makes this possibility less probable.

While the XRF signal cannot be quantified into absolute values, after corrections for the absorption cross-section, the relative abundance of metal ions can be compared to obtain the general trends in ion abundance. We therefore evaluated the atomic abundance of Zn, Mn, and Fe ions relative to the most abundant Ca ion in areas positioned at the biofilm bulk (away from the wrinkles) and compared them to the relative concentrations of these metal ions in the medium (Table S3).

We found that Ca accumulation is enhanced relative to the other metals ions, even after considering its higher abundance in the medium (Table S3). This suggests that calcium is preferentially bound in the biofilm thus maintaining a high concentration gradient relative to the medium. Ca binding by the ECM was already shown before, and suggested to stabilize the biofilm and inhibit biofilm dispersal (59). In contrast, Zn, Mn, and Fe ions are more free to diffuse and flow as solutes in water channels and they accumulate along wrinkles, where evaporation in enhanced (33). Strikingly, the relative abundance of Zn, Mn, and Fe with respect to Ca, is similar in ECM mutant biofilms and in the bulk of WT biofilms (away from wrinkles), and yet sporulation is detained. We suggest that even though the metal ions are accumulated across the film, the lack of wrinkles and lateral water flow in the mutant films prevents localized accumulation of some metal ions with respect to Ca, and that these altered Ca/metal ion ratios are crucial for sporulation. Interestingly, all these metal ions (Ca, Zn, Mn, and Fe) have been described to play a role in sporulation (59, 64, 65) and they also accumulated in isolated spore samples(Fig. S7).

To examine the possibility that Ca is bound to the matrix, we examined protein fibers of isolated TasA preparations with XRF. XRF analysis revealed that TasA fibers contained all the metal ions: Ca, Zn, Fe and in some cases Mn, that were also probed in WT and mutant biofilms (Fig. 5). However, different TasA polymorphs showed different preference to these metal ions as shown in an overlay of the XRF maps of these metal ions (Figs. 5D-5F and Fig. S8). In particular, in aggregates that formed in an acidic environment, iron binding dominated over Ca, Mn and Zn (Fig. 5D), whereas bundles that formed in high salt (NaCl) and protein concentrations accumulated more Ca relative to Zn, Mn and Fe (Fig. 5F, S8). The fibrils that formed at high salt concentration were especially heterogeneous, as was reflected both in different morphologies and in metal ion binding preferences residing in the same sample (Fig. 5E). These results suggest that some part of the metal ions in biofilms, especially Ca, are bound by TasA within the matrix. The variations in the XRF signal of TasA fibers, prepared under three different environmental conditions, is a further testimony of the protein’s polymorphism and its possible contribution to biofilm heterogeneity.

**Figure 5.**
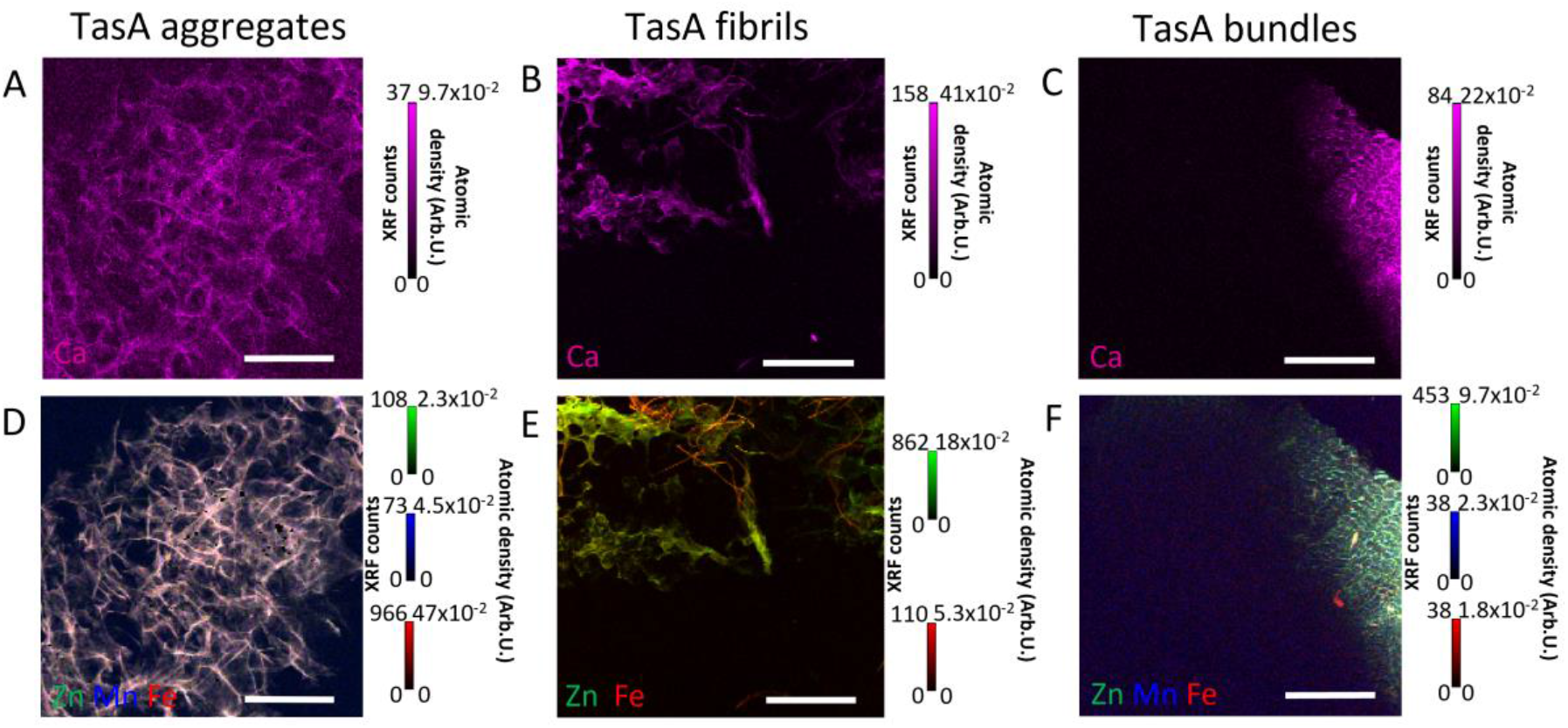
Morphology, metal ion binding and XRD profiles of TasA fibers formed in different environments. (A-F) XRF of TasA fibers, formed in acidic conditions (‘TasA aggregates), high salt concentration (‘TasA fibrils’) and high protein and salt concentration (‘TasA bundles’). XRF maps of Ca (A-C) are presented separately from the overlay of the XRF maps of Zn, Fe, and Mn (D-F) due to its higher abundance (atomic density) in all the samples. Color code: Ca (magenta), Zn (green), Mn (blue) and Fe (red); red and green = yellow. Scale bar, 300 μm.

## Discussion

Spatio-temporal analysis of molecular structures within *B. subtilis* biofilms reveals that cellular spatial differentiation by function also translates into structural and elemental heterogeneity across whole biofilms at the molecular level. Probing intact biofilms with XRD, we have found that *B. subtilis* biofilms convey three characteristic structural signatures originating from spores, ECM components, and water in bound and free state. Concomitant spatial elemental analysis by XRF showed that the metal ions, mainly, Ca, Zn, Mn, and Fe, known to play a crucial role in bacterial metabolism and sporulation (66), differentially accumulate in *B. subtilis* biofilms. While Ca is evenly distributed across the biofilms, other metal ions, mainly Zn, Mn and Fe, accumulate along biofilm wrinkles.

Based on our observations we suggest an inclusive view of biofilm development, linking between macroscopic features, namely biofilm wrinkles, to elemental and structural heterogeneities in biofilms, as illustrated schematically in Fig. 6. We suggest a dual role for the ECM in biofilms. It provides structural support that lead to wrinkle formation and it also serves as filter, selectively binding Ca over other metal ions. These two properties lead to heterogeneous distribution of metal ions and spores, as detailed below.

**Figure 6.**
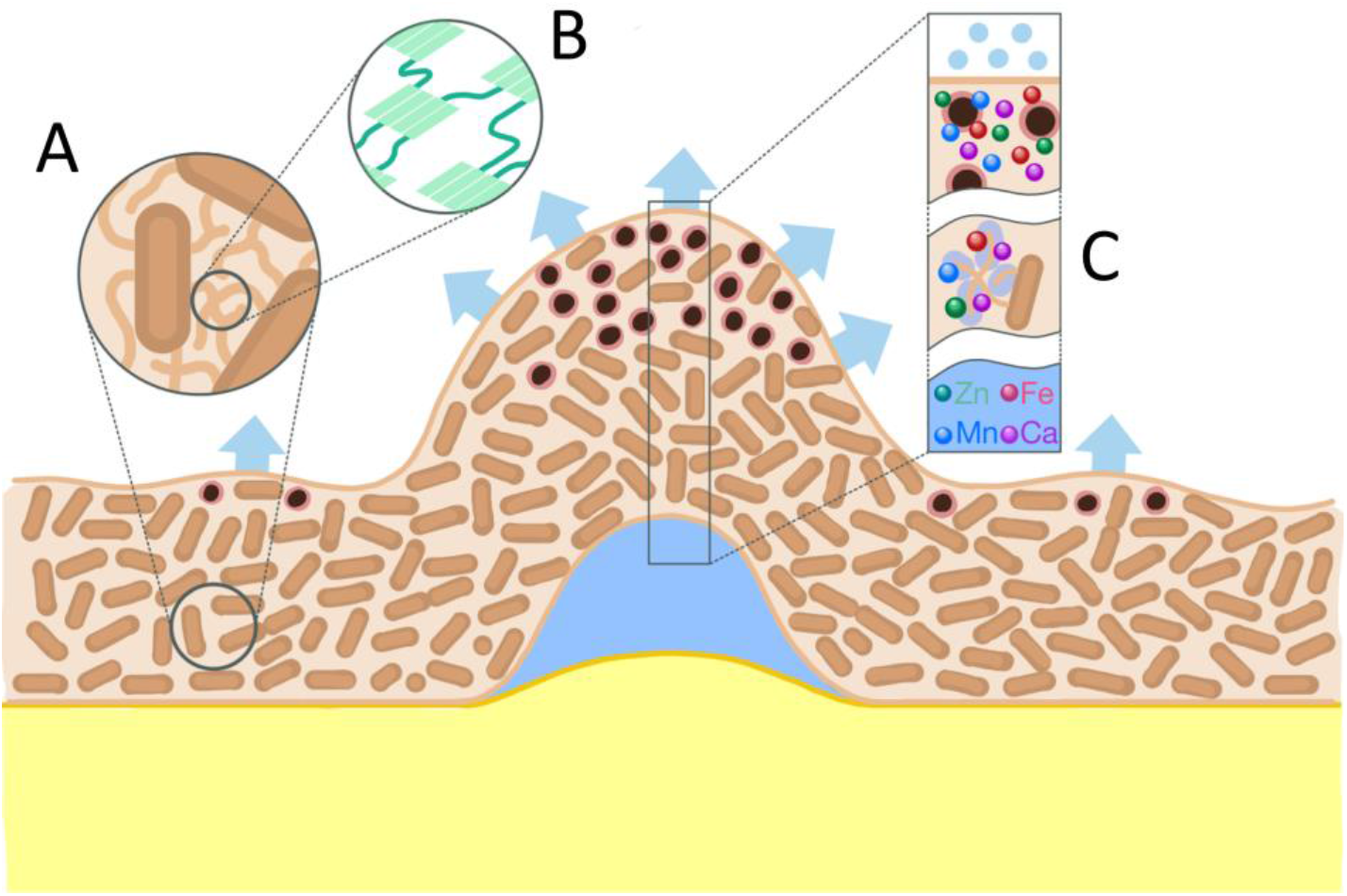
Schematic representation of the relationships between biofilm structures and metal ion distribution and their implications on biofilm physiology. The cartoon shows a side view of a WT *B. subtilis* biofilm, drawn around a wrinkle, and it captures the major findings of this study. We identified three dominant structure-based subgroups in biofilms, vegetative bacterial cells (represented by brown ovals surrounded by an orange coat), spores (represented by dark circles surrounded by a pink coat) extracellular matrix components (represented by a brown mesh surrounding the cells), and a water channel (shown in light blue) residing beneath a biofilm wrinkle. The light blue arrows represent water evaporation. A zoom into the ECM mesh shows ECM fibrillar components (A), and a further zoom in highlights their structure, short cross β sheet domains that are connected with non-cross β sheet regions (B). A column across the biofilm wrinkle (black rectangle) marks the three areas along the biofilm where metal ions reside (we only refer to the metal ions that were observed in this study). Water molecules and metal ions are free inside water channels. They are then adsorbed by the ECM, with Ca being more abundant than Zn, Mn, and Fe. Ca remains mostly bound to the ECM, but Zn, Mn, and Fe are free to diffuse and concentrated in biofilm wrinkles, where water evaporation is the largest, possibly leading to sporulation.

Metal ions, initially residing in the medium, accumulate in the ECM biopolymers with preferred Ca binding. This process is driven by water evaporation which occurs throughout the film (blue arrows in Fig. 6). However, Wilking et al., (33) have shown that biofilm wrinkles act as water channels, where water flow is driven by enhanced evaporation along the wrinkles. The water flow carries nutrients, but as Ca is selectively bound by the matrix and therefore filtered out, the solution is relatively enriched with Zn, Mn and Fe, which are only weakly bound by the matrix or taken up by cells. These ions eventually accumulate along the wrinkle due to water evaporation. The co-occurrence of these metal ions and spores along wrinkles is consistent with their essential role in sporulation (64, 67, 68) and points at the possibility that spores act as drainage source to metal ions as a means to circumvent toxicity. Our study also suggests that in the absence of wrinkles in ECM mutant biofilms, a Ca/metal ion ratio required for sporulation cannot be achieved, providing reason to the detained sporulation in ECM mutant biofilms.

Our model therefore offers a functional link between extracellular matrix properties, macroscopic wrinkles and sporulation via heterogenic metal ions distribution, showing that biofilm heterogeneity is not only affected by genetic expression and cellular differentiation, but also by the passive effects that stem from physicochemical properties of molecules secreted by the cells in the biofilms. These lead to differential distribution of nutrients that propagates through macroscopic length scales. Multiscale approaches to biofilm internal structures and metal ion relationships may hold key to understanding biofilm physiology and multicellularity and their relation to subpopulation survival in B. subtilis as well as in other biofilms of single or mixed bacterial species.

## Materials and Methods

### Bacterial *Bacillus subtilis* strains used in this study

Wild-type (NCIB361O)

eps mutant SSB488 (361O *epsA-O::tet*)

*tasA* mutant CA017 (361O *tasA::kan*)

sinReps double mutant (ZK4363)

### Biofilm sample preparation

A 2 μl drop of bacterial cell suspension, grown as liquid culture overnight at 37 °C in Luria-Bertani (LB) medium, was spotted on solid MSgg medium agar (1.5%) plate (69), dried and kept at 30 °C or room temperature between 12 h – 10 days, as indicated in the text.

### Spores sample preparation

Wild-type *Bacillus subtilis* LB liquid culture, grown for 8 h at 37 °C, was spread (200 μl suspension) on solid LB agar (1.5%) plate and kept for 48 h at 37 °C.

### TasA Purification and sample preparation

TasA was purified as described previously (34, 36). Briefly, sinR eps double mutant *B*.*subtilis* cells (ZK4363) were grown in MSgg broth, at a 1:100 dilution from an LB overnight culture, under shaking conditions. After 16 h of growth at 37 °C the cells were pelleted (10000 g, 15 min, 4 °C), resuspended with saline extraction buffer and probe sonicated (1518 J, 5 s pulse and 2 s pulse off). Following an additional centrifugation step (10000 g, 15 min, 4 °C), the supernatant was then collected and filtered through a 0.45 μm polyethersulfone (PES) bottle-top filter. The filtered supernatant was collected after centrifugation (17000 g, 15 min, 4 °C), concentrated with Amicon centrifugal filter tubes and either incubated in saline extraction buffer for 72 h at 4 °C (used to prepare ‘TasA bundles’) or passed through a HiLoad 26/60 Superdex S200 sizing column that was pre-equilibrated with a 50 mM NaCl, 5 mM HEPES solution at pH 8 (used to prepare ‘TasA aggregates’ and ‘TasA fibrils’). Purified proteins were stored at −20 °C until use. The concentration of the protein was determined using a BCA protein assay (ThermoFisher Scientific, Waltham, MA, USA).

### Polysaccharide purification

Polysaccharide was purified from WT Bacillus Subtilis pellicles, as previously described (38, 70). WT liquid culture was grown overnight in LB medium and transferred to SYN medium at a 1:100 dilution. Pellicles were collected after 24 h at 37 °C, transferred into fresh SYN medium and washed twice by centrifugation at 5000 rpm for 15 min at 25 °C. The pellet was collected probe sonicated (1518 J, 5 s pulse and 2 s pulse off) (Sonics, Vibra cell CV334) in fresh medium for 150 s. The supernatant was separated from the pellet after an additional sonication and centrifugation of 5000 rpm for 30 min at 25 °C. The EPS was precipitated by adding x3 ice-cold isopropanol (J.T. Baker) following an overnight at -20 °C. The supernatant was discarded and the precipitate was washed twice with ice-cold isopropanol and was kept at -20 °C for additional 2 h. The precipitant was collected, dissolved in 0.1 M MgCl_2_ (J.T. Baker) and extracted twice by phenol-chloroform (Fisher Chemical) on a separating funnel. The aqueous phase was collected, dialyzed for 48 h against DDW (with a Cellu Sep T3, 12000-14000 MWCO.) and lyophilized to dryness. For purification from DNA and proteins, EPS was dissolved in a 10 mM Tris buffer (VWR) containing 50 mM NaCl, 10 mM MgCl_2_ pH 7.5 and incubated with DNase (Sigma-Aldrich) at 37 °C for 1 h. Proteinase K (Sigma-Aldrich) was then added and the solution was incubated at 40 °C for 1 h. The solution is dialyzed against DDW (with a Cellu Sep T3, 12000-14000 MWCO.) for 48 h and lyophilized to dryness.

### TasA fibers preparation

TasA fibers were prepared as previously described (53), under the following conditions:

TasA bundles: Purified TasA was incubated in saline extraction buffer for 72 h at 4 °C. The solution was then dialyzed against TDW for three days, lyophilized and placed between two silicon nitride membrane or resuspended in a drop of water and mounted on a loop (Hampton Research).

TasA fibrils: Purified TasA was passed through a HiLoad 26/60 Superdex S200 sizing column that was pre-equilibrated with a 50 mM NaCl and 5 mM HEPES solution at pH 8. Sodium chloride stock solution (5M) was then added to the protein yielding the final concentration, 1.5 M NaCl and 130 μg/ml TasA. The solution was incubated for 72 h at 4 °C and dialyzed against TDW for three days, lyophilized and placed between two a silicon nitride membrane.

TasA aggregates: Purified TasA was passed through a HiLoad 26/60 Superdex S200 sizing column that was pre-equilibrated with a 50 mM NaCl and 5 mM HEPES solution at pH 8. The solution pH was then adjusted to 2.5 with formic acid, yielding TasA aggregates (52) and kept at room temperature for three days at 4 °C. The solution was exchanged by consecutive centrifugation (5000 rpm, 10 min) and supernatant replacement with TDW, three times every 24 h. The solution was then lyophilized and placed between two silicon nitride membranes or resuspended in a drop of water and mounted on a loop (Hampton Research).

### Sample preparation for XRD measurements

Biofilm samples were mounted on aluminium frames overlaid with kapton films (13 μm thick from Goodfellow Cambridge limited or 25 μm thick from DuPont) or on a silicon nitride membrane (Silson Ltd., frame: 10.0 mm × 10.0 mm, 200 μm thick; membrane: 5.0 mm × 5.0 mm, 1000 nm thick). Hydrated biofilm measured at myspot was gently pulled off the agar plate, and sealed at the lab one hour prior to the measurement between two kapton films. During biofilm removal, only WT biofilms maintained their shape and humidity level. Fixed samples were prepared by overnight exposure to 15% formaldehyde vapour following mounting on kapton foil/Silicon nitride membrane.

Spore sample was scrapped from the plate and placed between Kapton sheets that were glued on an aluminium frame and sealed with a parafilm. Protein samples measured at ESRF were sandwiched between two 1000 nm thick silicon nitride membranes (see above).

### SAXS/XRD and XRF mapping

Scanning XRD/XRF measurements of biofilms and proteins were performed at the mySpot beamline synchrotron BESSY II (Helmholtz Center, Berlin, Germany) and at the ID13 beamline Synchrotron ESRF (Grenoble, France).

At ID13 (ESRF) the X-ray beam was defined to 13.0 keV (0.954 Å) using Si(111) channel-cut monochromator and focused to 2.5 μm cross section. Scan steps varied between 2 μm or 5 μm. 2D patterns were acquired in transmission mode using a Dectris, EIGER X 4M detector (pixel size 0.075×0.075 mm^2^), by integrating for 2 s in an “on-the-fly” scanning mode. A Vortex EM, silicon drift detector with 50 mm^2^ sensitive area, XRF detector was placed roughly 90°to the incoming X-ray beam to collect the emitted X-ray photons from the sample simultaneously. Acquisition time was set to 2 s/point.

At mySpot beamline the X-ray beam (Bessy II) was defined to 15.0 keV (0.827Å) using a B4C/Mo Multilayer monochromator and focused to (50×50) μm^2^ using KB mirrors and 50 μm^2^ pinhole. XRD data were obtained using a 2D Dectris Eiger 9 M detector ((2070 × 2167) pixel^2^) in transmission geometry, and a single element Si(Li) XRF detector (RAYSPEC (SGX Sensortech) was placed perpendicular to the beam. Acquisition time was set to 60 s/point.

Sample to detector distance (around 300 mm) was calibrated using quartz powder in both setups. When present, internal kapton signal at *q* = 4 nm^-^1 was used to further verify the calibration for each sample. Calibration of XRF detectors was performed using pyMCA (http://pymca.sourceforge.net/) using the argon emission and the main beam scattering energy. XRF intensities were corrected by the theoretical absorption cross-section for each element, at 15 or 13 keV (60-63) taking into account (when relevant) the absorption of the XRF by the detector Beryllium window (8 μm thick).

### SAXS/WAXS/XRF analysis

Calibration, integration background removal, and peak-fitting of the 2D diffraction patterns were performed using the software DPDAK (71) or home-written routines based on pyFAI (https://pyfai.readthedocs.io/en/master/). The fitted parameters as well as the integrated XRF signal around specific emission lines, were exported and plotted in Origin 2020. For 1D presentation, the 1D patterns from specific regions of interest (ROI)s (X, Y, Z in Fig. 2F) were averaged and normalized by subtracting the minimum value and dividing by the maximum intensity.

The XRF maps were analysed using the software ImageJ (72). Average atomic densities were extracted by analysing the calibrated grey-level histograms of relevant ROIs in the maps. The correction of the fluorescence intensity signal to relative atomic density was performed taking into account the absorption cross section at E=15 keV (Bessy II data), 13 keV (ESRF data) (60-63). We note that absolute values cannot be quantitatively compared between beamline set ups and between sample as they depend amongst others on the incoming flux, the sample to detector distance and exact geometry, and the sample thickness.

### Transmission Electron Microscopy of Biofilm colonies

Agar slabs (less than 0.5 mm^2^) containing defined regions of the bacterial colonies were cut out and inverted on lecithin-coated 3-mm specimen carriers for high pressure freezing in a high-pressure freezing machine (Leica, ICE) with or without 10% polyvinylpyrrolidone as a filler (Δ*eps* biofilms and one of the WT samples were frozen without filler). Samples were transferred under liquid nitrogen into a freeze substitution unit (Leica, AFSII) with 0.1% osmium tetroxide in acetone for substitution medium. Samples were slowly warmed from -90 °C to 0 °C within 3 days.

The samples were washed with acetone for one hour and incubated with 0.5% uranyl acetate in acetone for 1 h still at 0 °C. Samples were washed 3x with acetone (moving from 0 °C to room temperature), before they were infiltrated with 25% and 50% epoxy resin (Roth, Germany) in acetone for 90 min. Samples were left in 75% epoxy resin overnight. Four changes of 100% epoxy resin followed over the period of 2 days before samples were flat-embedded between Teflon-coated microscope slides.

Infiltrated samples were polymerized at 60 °C for two days and defined regions mounted on resin stubs for cutting 70 nm–ultrathin cross-sections through the bacterial colonies, which were post-contrasted with 0.5 % uranyl acetate and 0.04% lead-citrate. Samples were investigated on a Morgagni transmission electron microscope (Thermofisher/FEI/Philips) at 80 kV and micrographs acquired with a Morada CCD camera (EmSis/Olympus) with ITEM software (EmSis/Olympus/Sys). Images were analysed using Fiji software (72)).

### TEM images analysis according to cell type

Unbiased micrographs (image area (6 × 4.7) um with a 1.87 nm/px resolution) of the biofilm samples were acquired using the xy-coordinate display at the TEM, with a step size of 20 μm. Cells were marked and counted in these TEM images (N, total number of cells = 7159). Cell types were classified as normal cells, lysed cells, deformed cells, sporulating cells or spores as indicated in the text and in Fig. S6. The number of cells of each type was normalized by the number of images evaluated, which normalizes the total area that was probed. Normalizing by the area takes into account differences in cellular density and therefore we chose to normalize by area and not by total cell number.

## Acknowledgments

*Measurements were carried out at the mySpot beamline at the BESSY II electron storage ring operated by the Helmholtz-Zentrum Berlin für Materialien und Energie and at beamline ID13 at the European Synchrotron Radiation Facility. We thank the HBZ and the ESRF for provision of synchrotron radiation facilities and for financial support*. We thank EM facility of MPI-CBG for support, use of equipment and reagents, Daniel Werner at the MPIKG for X-ray measurements. Jiliang Liu for ESRF ID13 beamline assistance at ESRF. Special thanks to Prof. Sigal Ben Yehuda for insightful discussions, Yosef Edery Aharony for his contribution with data sorting and image analysis and Daniel Rosenblatt for assistance with the image analysis. This work was funded by the Kaete Klausner scholarship.

