## Supporting information for "Multiscale X-ray study of *Bacillus subtilis* biofilms reveals interlinked structural hierarchy and elemental heterogeneity"

### equal contribution

**Table S1. Fitting results - Peak fitting results. Corresponding fits are plotted in Figs. S1, S4.**

| | Low range | | | | | | | | | High range | | | | | | | | | | | |
|---|---|---|---|---|---|---|---|---|---|---|---|---|---|---|---|---|---|---|---|---|---|
| **Samples** | **Hump** | | | **Peak i** | | | **Peak ii** | | | **Hump** | | | | | | | | | | | |
| | Position q ($nm^{-1}$) | Width q ($nm^{-1}$) | Intensity (arb. U.) | Position q ($nm^{-1}$) | Width q ($nm^{-1}$) | Intensity (arb. U.) | Position q ($nm^{-1}$) | Width q ($nm^{-1}$) | Intensity (arb. U.) | Position q ($nm^{-1}$) | Width q ($nm^{-1}$) | Intensity (arb. U.) | Position q ($nm^{-1}$) | Width q ($nm^{-1}$) | Intensity (arb. U.) | Position q ($nm^{-1}$) | Width q ($nm^{-1}$) | Intensity (arb. U.) | Position q ($nm^{-1}$) | Width q ($nm^{-1}$) | Intensity (arb. U.) |
| WT 72 | 6.52 | 1.22 | 0.67 | 6.29 | 0.13 | 0.04 | 6.75 | 0.09 | 0.03 | 13.58 | 1.00 | 0.54 | 14.58 | 2.64 | 4.09 | 18.00 | 3.83 | 2.37 | - | - | - |
| WT hydrated wrinkle | 5.4 | 1.49 | 0.39 | 6.2 | 0.07 | 0.02 | 6.63 | 0.08 | 0.03 | 13.81 | 1.65 | 0.44 | - | - | - | 19.43 | 2.63 | 4.44 | 25.9 | 6.22 | 6.5 |
| WT Hydrated bulk | 5.4 | 1.41 | 0.75 | 6.2 | 0.06 | 0.02 | 6.63 | 0.07 | 0.03 | - | - | - | 14.64 | 2.29 | 0.75 | 19.19 | 1.25 | 0.68 | 19.95 | 2.83 | 5.23 |
| WT 24 | 6.16 | 1.19 | 0.56 | - | - | - | - | - | - | 13.82 | 1.63 | 1.58 | 15.76 | 3.06 | 3.92 | 18.00 | 11.64 | 6.50 | - | - | - |
| WT12 with hump | 6.26 | 1.28 | 0.57 | - | - | - | - | - | - | 13.72 | 1.75 | 2.11 | 16.22 | 3.19 | 4.10 | - | - | - | 23.49 | 7.02 | 2.48 |
| WT12 no hump | - | - | - | - | - | - | - | - | - | 13.94 | 2.24 | 4.26 | - | - | - | 18.57 | 2.77 | 3.30 | 27.00 | 5.46 | 6.50 |
| ΔEPS with doublet | 6.25 | 1.42 | 1.02 | 6.24 | 0.15 | 0.02 | 6.71 | 0.14 | 0.02 | 13.66 | 1.14 | 0.59 | 15.51 | 2.86 | 3.68 | 18.00 | 6.80 | 5.85 | - | - | - |
| ΔEPS without doublet | 6.00 | 1.17 | 0.82 | - | - | - | - | - | - | 13.43 | 1.27 | 1.06 | 15.25 | 2.38 | 3.37 | 18.99 | 3.37 | 1.93 | - | - | - |
| ΔTasA | 6.23 | 1.02 | 0.33 | - | - | - | - | - | - | 13.66 | 1.60 | 1.23 | 16.09 | 2.78 | 3.32 | 18.00 | 8.50 | 5.90 | - | - | - |
| ΔEPSΔTasA | 6.21 | 1.01 | 0.25 | - | - | - | - | - | - | 13.83 | 1.59 | 1.39 | 15.67 | 3.18 | 4.52 | 18.00 | 6.74 | 3.42 | - | - | - |
| Spores | 6.27 | 1.85 | 2 | 6.25 | 0.15 | 0.03 | 6.72 | 0.15 | 0.03 | 13.58 | 0.63 | 0.23 | 14.36 | 2.78 | 5.56 | 18.00 | 4.34 | 3.92 | - | - | - |
| EPS powder | 7.50 | 1.49 | 0.54 | - | - | - | - | - | - | 12.27 | 2.10 | 4.77 | - | - | - | 17.28 | 1.86 | 1.23 | 23.09 | 7.43 | 6.49 |
| TasA aggregates | 6.48 | 1.43 | 0.85 | - | - | - | - | - | - | 13.77 | 1.14 | 0.83 | 14.75 | 3.44 | 5.88 | 18.00 | 8.81 | 2.69 | - | - | - |
| TasA bundles | 6.51 | 1.15 | 0.78 | - | - | - | - | - | - | 13.90 | 1.32 | 1.04 | 15.73 | 2.82 | 4.10 | 18.00 | 11.62 | 6.5 | - | - | - |
| TasA fibrils | 6.25 | 1.27 | 0.48 | - | - | - | - | - | - | 13.93 | 1.33 | 1.01 | 16.16 | 4.19 | 6.09 | - | - | - | 26.82 | 2.81 | 0.79 |
| Water | - | - | - | - | - | - | - | - | - | - | - | - | - | - | - | 19.52 | 2.68 | 4.38 | 26.33 | 5.52 | 6.5 |

**Table S2. Fitting results - real space *d*-spacing**

| | Low range | | | | | | High range | | | |
|---|---|---|---|---|---|---|---|---|---|---|
| Sample | Hump / d (Å) | | Peak I / d (Å) | | Peak ii / d (Å) | | Hump / d (Å) | | | |
| | Position | Width | Position | Width | Position | Width | Position | Width | Position | Width |
| WT 72 | 9.63 | 1.81 | 9.99 | 0.21 | 9.30 | 0.12 | 4.63 | 0.34 | 4.31 | 0.78 |
| WT hydrated wrinkle | 11.64 | 3.21 | - | - | - | - | 4.55 | 0.54 | - | - |
| WT Hydrated bulk | 11.64 | 3.04 | - | - | - | - | - | - | 4.29 | 0.67 |
| WT 24 | 10.19 | 1.97 | - | - | - | - | 4.55 | 0.54 | 3.99 | 0.78 |
| WT12 with hump | 10.04 | 2.05 | - | - | - | - | 4.58 | 0.59 | 3.87 | 0.76 |
| WT12 no hump | - | - | - | - | - | - | 4.51 | 0.72 | - | - |
| ΔEPS with doublet | 10.05 | 2.29 | 10.06 | 0.24 | 9.36 | 0.20 | 4.60 | 0.38 | 4.05 | 0.75 |
| ΔEPS without doublet | 10.47 | 2.04 | - | - | - | - | 4.68 | 0.44 | 4.12 | 0.64 |
| ΔTasA | 10.08 | 1.64 | - | - | - | - | 4.60 | 0.54 | 3.91 | 0.68 |
| ΔEPSΔTasA | 10.12 | 1.64 | - | - | - | - | 4.54 | 0.52 | 4.01 | 0.81 |
| Spores | 10.03 | 2.97 | 10.06 | 0.24 | 9.36 | 0.20 | 4.63 | 0.21 | 4.38 | 0.85 |
| EPS powder | 8.38 | 1.66 | - | - | - | - | 5.12 | 0.88 | - | - |
| TasA aggregates | 9.70 | 2.14 | - | - | - | - | 4.56 | 0.38 | 4.26 | 0.99 |
| TasA bundles | 9.65 | 1.70 | - | - | - | - | 4.52 | 0.43 | 3.99 | 0.72 |
| TasA fibrils | 10.05 | 2.03 | - | - | - | - | 4.51 | 0.43 | 3.89 | 1.01 |

**Table S3.** **Ca/metal ratios of the relative atomic density of Ca to the metals: Mn, Zn, Fe in biofilms calculated based on the XRF intensity and corrected for the atomic absorption cross-sections at energy of 13 keV. Note that the ratios only represent a qualitative estimate of the relative abundance of the different metal ions. Quantitative comparison between samples and across beam-time sessions is hindered by variations in sample thickness and morphology and the precise measurement geometry and beam flux. Corrected ratios denote the Ca/metal ratios following division in the same ratios in the medium. WT and mutants were measured in different beamtime sessions.**

| XRF | Ca/metal ratios | | | | | |
|---|---|---|---|---|---|---|
| Sample | Ratios | | | Ratios corrected with medium | | |
| | Ca/Mn | Ca/Zn | Ca/Fe | Ca/Mn | Ca/Zn | Ca/Fe |
| **WT 72 (bulk)** | 6.1 ± 0.6 | 110 ± 10 | 13.3 ± 0.4 | 3 ± 1 | 3 ± 2 | 1.3 ± 0.8 |
| **WT 24** | 10.4 ± 0.3 | 104 ± 5 | 13.4 ± 0.2 | 6 ± 2 | 2 ± 2 | 1.3 ± 0.9 |
| **WT 12** | 16 ± 4 | 140 ± 60 | 18 ± 3 | 9 ± 4 | 3 ± 3 | 2 ± 1 |
| **ΔEPS** | 8.3 ± 0.7 | 101 ± 6 | 12.4 ± 0.1 | 5 ± 2 | 2 ± 2 | 1.2 ± 0.8 |
| **ΔTasA** | 8 ± 1 | 132 ± 15 | 15.4 ± 0.1 | 4 ± 2 | 3 ± 2 | 1 ± 1 |
| **ΔEPSΔTasA** | 12 ± 2 | 90 ± 30 | 12 ± 2 | 6 ± 3 | 2 ± 2 | 1.2 ± 0.8 |
| **MSgg/agar** | 1.8 ± 0.7 | 40 ± 30 | 10 ± 7 | | | |

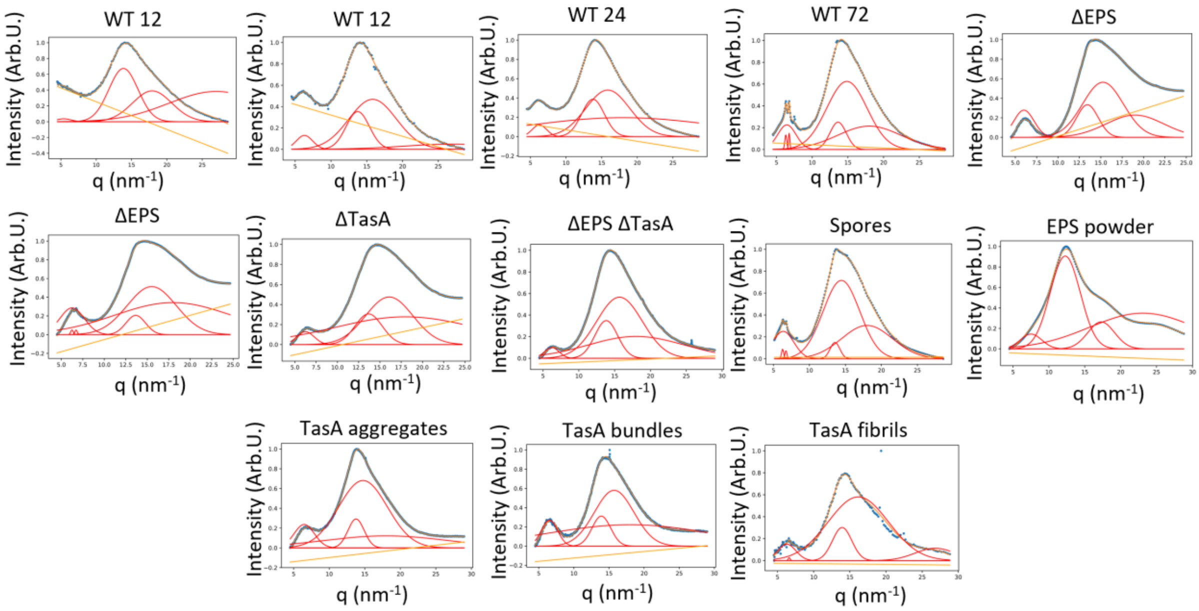

**Fig. S1**. Peak fitting results for WT and mutant biofilms, spores and TasA fibers. Data: blue dotted line, fitted line: orange curve, baseline: yellow, fitted peaks: red

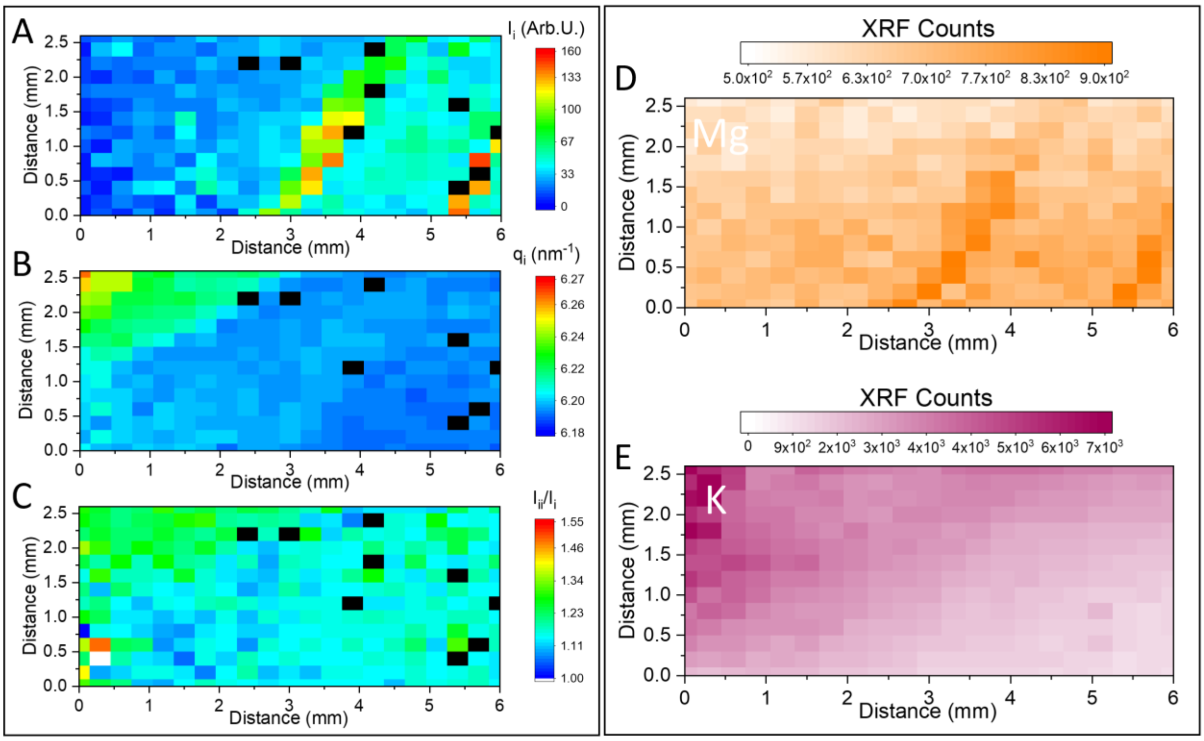

**Fig. S2**. XRD and XRF results of intact sealed 10 days old WT biofilm. (A) mapping the intensity of peak (i) showing an increase along the wrinkles pattern. (C) mapping the doublet peak intensity ratio (peak (ii)/peak (i)) showing an overall uniform value across the entire sample. (B) mapping of the position of the first doublet peak (peak (i) in Fig. 2C), showing a gradient shift with increasing hydration. (D), (E) XRF intensity of the magnesium and potassium signal, respectively. Mg and K XRF signals are not corrected for the absorption cross section of the elements.

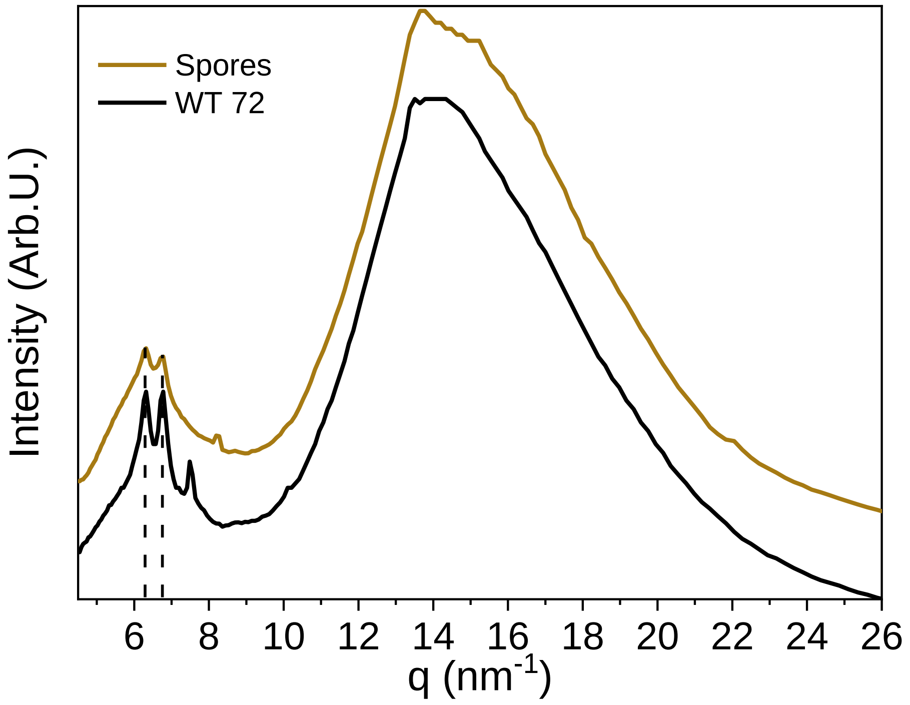

**Fig. S3.** 1D XRD profile of a fixed and dry 72 h old biofilm sample ('WT72', black line) overlayed with the 1D profile of a spore-containing sample ('Spores', brown curve).

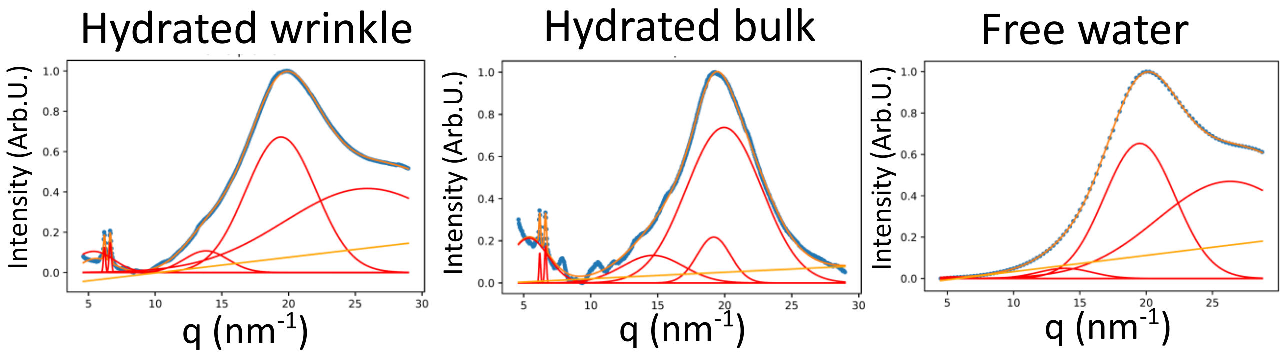

**Fig. S4**. Peak fitting results for WT biofilms that vary in hydration state and free water (see definitions in the main text and in Fig. 2C). Data: blue dotted line, fitted line: orange curve, baseline: yellow, fitted peaks: red.  The curve for Free water is taken from reference [44].

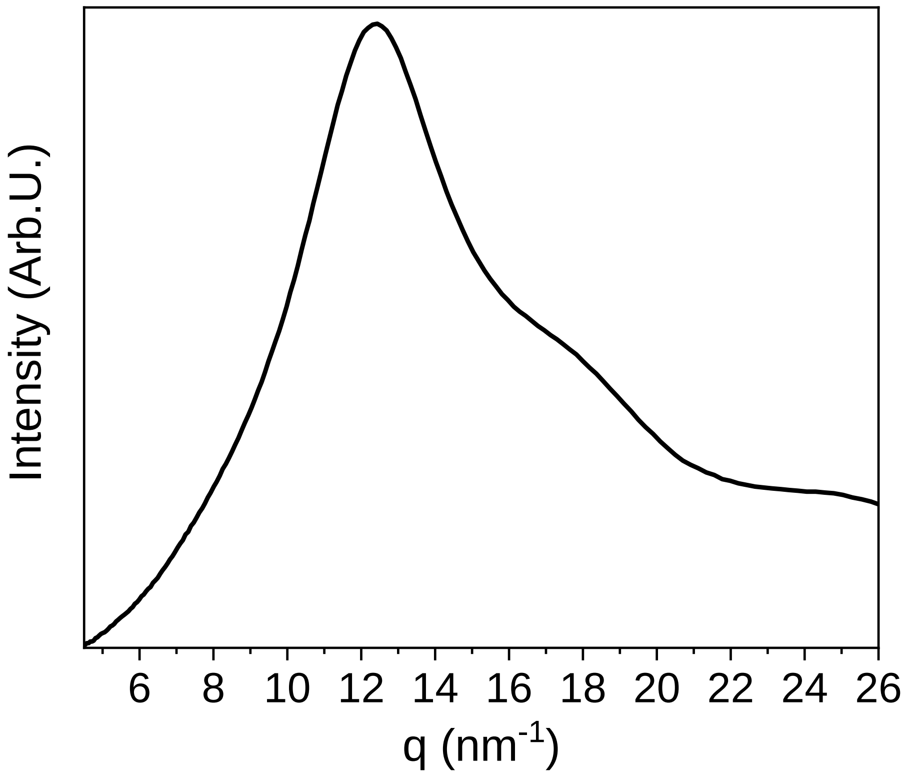

**Fig. S5**. 1D XRD profile of a polysaccharide, isolated from WT pellicles (refs. 38, 70).

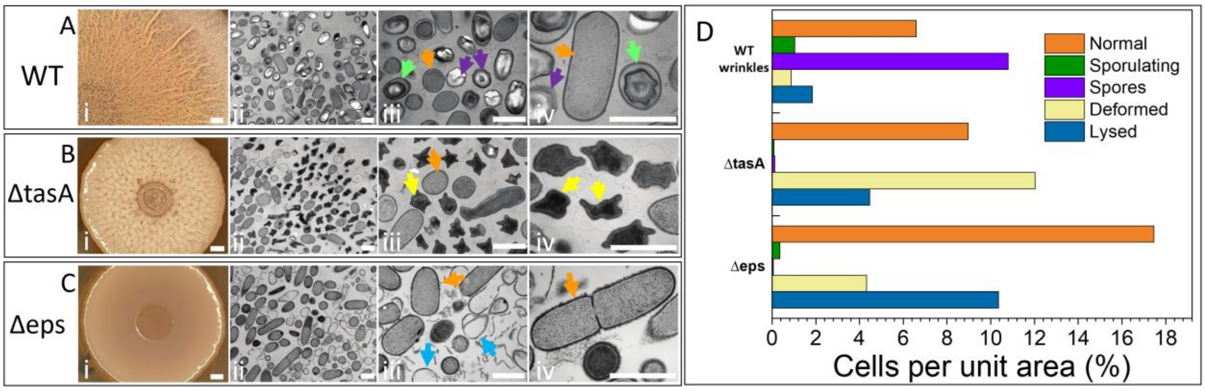

**Fig. S6.** Cell type abundance in WT and matrix mutant biofilms. ($A_i$, $B_i$, $C_i$) Light microscopy of biofilm samples and TEM imaging of fixed and stained thin sectioned WT ($A_{ii}$ - $A_{iv}$), Δ*tasA* ($B_{ii}$-$B_{iv}$) and Δ*eps* ($C_{ii}$-$C_{iv}$) mutant biofilms. We identified several cell types in the TEM images: normal cells (marked orange), lysed cells (marked blue), deformed cells (marked yellow), sporulating cells (marked green) and spores (marked purple). Scale bar in all panels is 1 µm. (D) Quantitative analysis of the cell types observed in TEM images, as shown in Figs. S6$A_{ii}$-S6$A_{iv}$, S6$B_{ii}$-S6$B_{iv}$, S6$C_{ii}$-S6$C_{iv}$(total number of cells was 4715 in a total of 189 images). Number of cells, normalized by unit area (unit area = 28.14 $mm^2$ is the area of a single image). Cell types were identified and numbered as explained in the text and in the Experimental section.

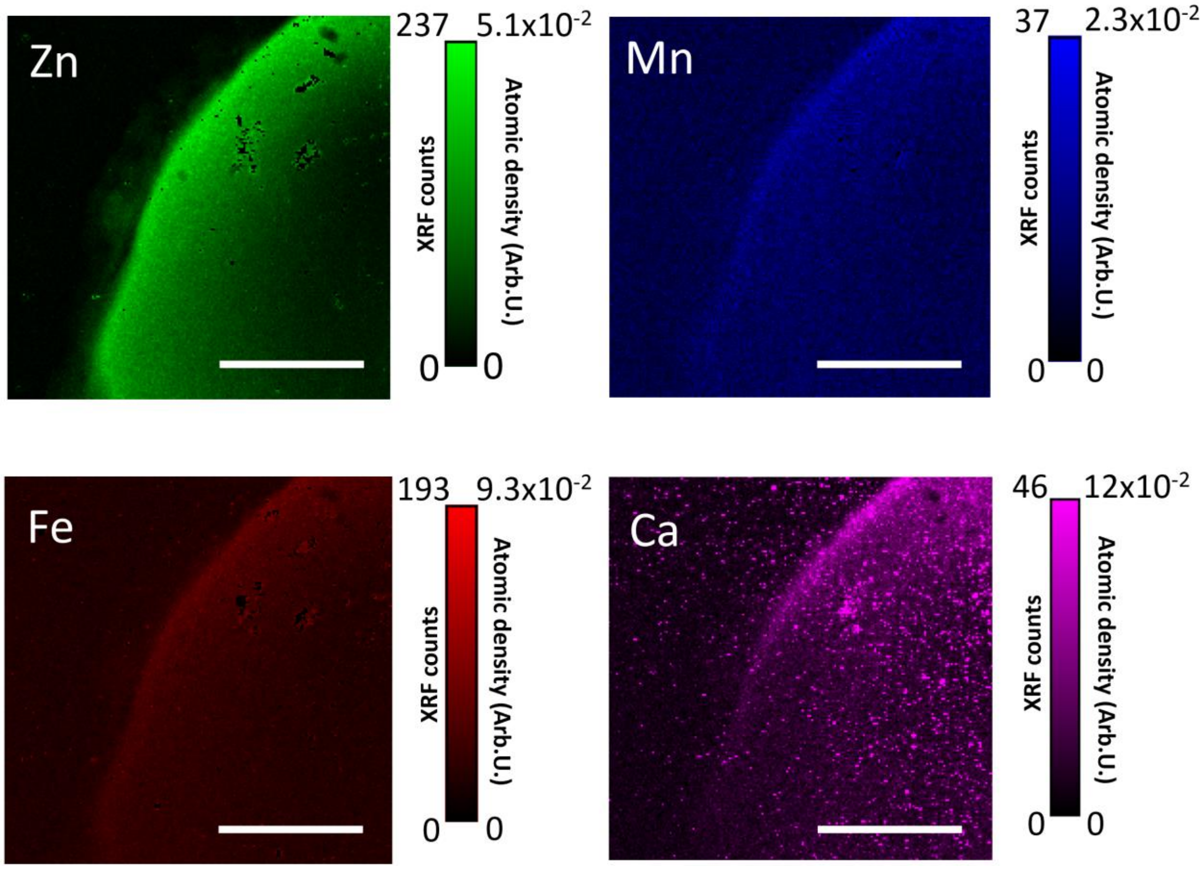

**Fig. S7.** XRF measurement of a spore - containing sample, The LUT shows the XRF counts and atomic density of Zn, Mn, Fe, and Ca. Salt precipitates were cut off the Zn, Mn and Fe XRF images, but they were kept in the Ca image to avoid cutting off the Ca signal. Scale bar represents 300 µm.

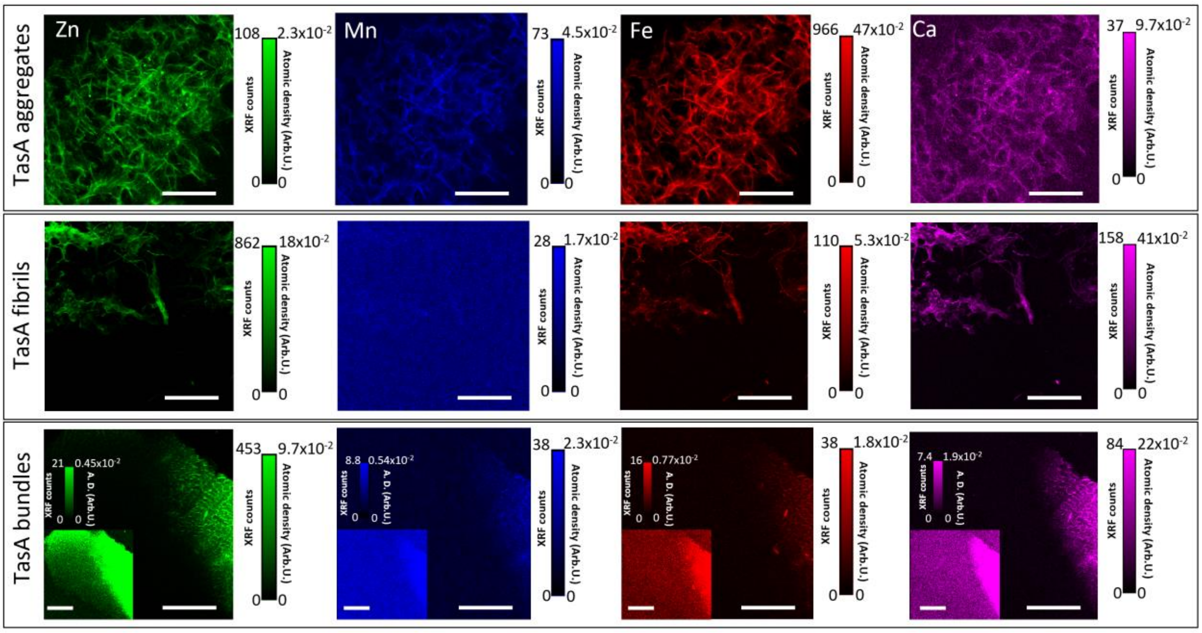

**Fig. S8**. XRF of TasA fibers formed in different conditions. Fibers formed in acidic conditions ('TasA aggregates), fibers formed in high salt concentration ('TasA fibrils') and fibers formed in high protein and salt concentration ('TasA bundles'). XRF of different metal ions is shown on separate panels. Color code: zinc (green), manganese (blue), iron (red) and calcium (magenta). The inset of the TasA bundles shows a low cut off intensity of the image, in order to emphasize low thickness areas in the sample. Scale bar represents 300 µm.

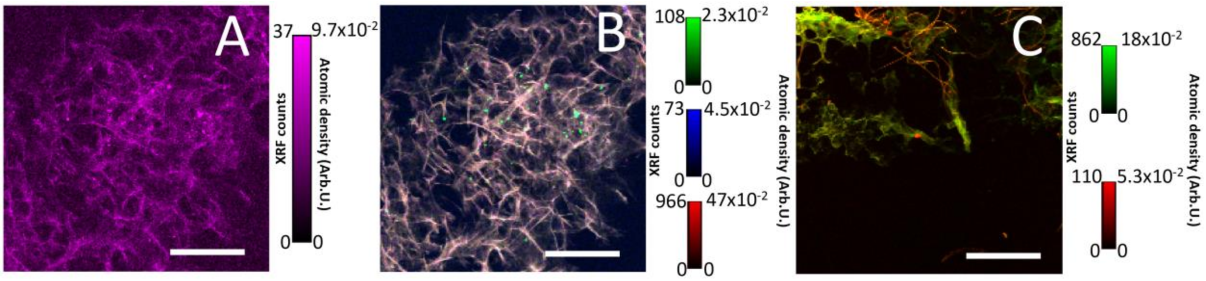

**Fig. S9**. Original XRF images of metal ions within TasA aggregates and fibrils. (A) XRF Ca map and (B) merge of Zn, Mn and Fe XRF of TasA aggregates. (C) Merge of Zn and Fe XRF of TasA fibrils. We present the XRF maps, shown in Fig. 5., without masking the higher signal of salt precipitates. Scale bar represents 300 μm.
